# Deep EEG Source Localization via EMD-based fMRI High Spatial Frequency

**DOI:** 10.1101/2022.11.29.518264

**Authors:** Narges Moradi, Bradley G. Goodyear, Roberto C. Sotero

## Abstract

Brain imaging with a high-spatiotemporal resolution is crucial for accurate brain-function mapping. Electroencephalography (EEG) and functional Magnetic Resonance Imaging (fMRI) are two popular neuroimaging modalities with complementary features that record brain function with high temporal and spatial resolution, respectively. One popular non-invasive way to obtain data with both high spatial and temporal resolutions is to combine the fMRI activation map and EEG data to improve the spatial resolution of the EEG source localization. However, using the whole fMRI map may cause spurious results for the EEG source localization, especially for deep brain regions. Considering the head’s conductivity, deep regions’ sources with low activity are unlikely to be detected by the EEG electrodes at the scalp. In this study, we use fMRI’s high spatial-frequency component to identify the local high-intensity activations that are most likely to be captured by the EEG. The 3D Empirical Mode Decomposition (3D-EMD), a data-driven method, is used to decompose the fMRI map into its spatial-frequency components. Different validation measurements for EEG source localization show improved performance for the EEG inverse-modeling informed by the fMRI’s high-frequency spatial component compared to the fMRI-informed EEG source-localization methods. The level of improvement varies depending on the voxels’ intensity and their distribution. Our experimental results also support this conclusion.

## 1. Introduction

Deep brain structures are critically important for brain function [1–4]. Thus, detecting the sources generated by a activity involving deep regions is crucial for evaluating these regions’ functions. Invasive techniques based on subdural electrodes are used to spot deep brain activity [1, 2, 5]. However, as these techniques are invasive, they could cause infection and other serious problems. Thus, a non-invasive yet accurate method is desired for localizing deep brain activity.

Electroencephalography (EEG) noninvasively records the electric potential produced by synchronous electrical dipoles (generators or sources) in the brain. EEG signals are calculated by modeling the head layers with different conductivities and source currents using the quasi-static approximations of Maxwell’s equations. However, finding the sources creating the scalp EEG signal is quite challenging and suffers from imprecision (typically in the order of centimeters) [6–9]. It is due to the EEG inverse problems being mathematically ill-conditioned and ill-posed. In addition to needing precise modeling of sources’ positions and orientations, the head’s conductivity, and noises, the number of possible sources outnumbers the EEG electrodes on the scalp. Consequently, EEG inverse problems’ solutions are unstable due to their high sensitivity to measurement noise, and there could be many source configurations that produce the same electric potential on the scalp [1, 9–14].

A variety of methods has been proposed to solve the inverse problem by adding information and constraints on the sources and limiting the possible solutions [1, 6, 7, 9, 12–17]. Depending on the assumptions about the sources inside the head, EEG inverse modeling is categorized as parametric (dipole source localization and spatial filtering methods) and nonparametric (distributed dipole model) methods. Parametric models constrain the active sources to a limited number of focal brain sources and often fail to localize sources of activities involving widespread brain areas. On the contrary, nonparametric methods consider sources over all the brain volume or cortical surface [1, 9, 12–17].

However, the nonparametric EEG source-localization methods tend to assign sources of the recorded EEG signals to the cortical surface and near the sensors [1, 10, 18, 19]. Therefore, localizing EEG signals arising from deep brain sources is usually more complex and still under debate [1, 20]. Some methods, such as weighted Minimum Norm Estimation (wMNE) [21–24] and Low Resolution Brain Electromagnetic Tomography (LORETA) [25–27], have been proposed to reduce the bias toward surface sources and improve the accuracy of specifying active sources in deep brain locations such as sources in the thalamus and hippocampus [16, 26]. wMNE method uses the column normalization of the lead-field matrix as a weighting matrix to compensate for the lower gains of deeper sources in the MNE method. On the other hand, LORETA assumes similarities between the current density of each cortex point to its neighbors and, thus maximum smoothness of the solution. In the LORETA method, a Laplacian operator is combined with the lead-field normalization of the wMNE method to ensure spatial coherence [1, 16, 24, 26]. However, each of these methods has its disadvantages. For instance, LORETA localizes deep sources smoother and better compared to the wMNE technique but provides lower spatial resolution, which is not good for the focal source estimation [1, 16, 19, 20, 28].

A popular way to improve the estimation of deep activities is to guide EEG source localization by functional Magnetic Resonance Imaging (fMRI) data which is a noninvasive brain-imaging method with high spatial resolution. Combining high-spatial-resolution data from fMRI with high-temporal-resolution data from EEG could leads to data with a high spatial and temporal resolution [6, 7, 9, 29, 30]. In these approaches, EEG source localization results are affected by adding fMRI activation map information as a weight for locations that are most likely to be active during a specific task and condition. Thus, the fMRI activation map is used as a spatial constraint to solve the EEG inverse problem at a selected time window to investigate the brain-function dynamics [6–8].

Considering the fMRI map as a constraint should be done cautiously due to the mismatch between EEG and fMRI signals caused by the spatial extent of the BOLD signals around neuronal firing areas and signal detection failure [7, 8, 31, 32]. EEG electrodes are placed on the scalp, and according to Poisson’s equation, the electric field decays with the inverse of the square distance between the source and the sensors. Moreover, due to the electrical conductivity of the head (transduction of the signals through the brain, cerebrospinal fluid, skull, and scalp), deep sources’ electrical activity must be higher than a threshold to have a chance to be recorded by the EEG electrodes [2, 3, 5, 33–35]. Consequently, EEG electrodes might not capture all the activity revealed on the fMRI map [2, 33–35]. Thus, considering the whole fMRI activation map as prior information on activated areas could cause spurious results for the EEG source localization.

Several methods have been proposed to enhance the integration of the fMRI activation map with the EEG inverse modeling. Some methods have used the Bayesian framework to probabilistically incorporate fMRI constraints into the EEG source localization. Bayesian-based models add fMRI activation information in the form of statistical priors and try to find the most appropriate combination of priors to be added to the EEG inverse modeling [7, 29, 36, 37].

In this paper, we show that using fMRI’s high spatial frequency components instead of the whole fMRI map provides more detailed spatial information on places with the local high activation [38]. It has been shown that, in active areas, lower activation intensity and, accordingly, lower signal-to-noise ratio (SNR) around the highly activated voxels may reflect the spatial extent of the BOLD signals around neuronal firing spots [32]. Thus, fMRI high spatial frequency component-by specifying the local high-intensity voxels, representing neuronal electrical activity, in active areas-spots the neuronal activities that are more likely to be recorded by the EEG electrodes. Moreover, the highly activated voxels on the fMRI map are more likely to have enough SNR to be captured by the EEG electrodes [2, 33–35] and the activation intensity threshold should be applied to these specified local highly activated voxels.

A fMRI activation map is generated by the general linear model (GLM), considering conditions as regressors for the task-based data [2, 39]. We use the three-dimensional EMD method [40–42] to decompose each fMRI activation map into its 3D or Spatial Intrinsic Mode Functions (3D-IMFs or SIMFs). EMD is an adaptive and data-driven method that applies to any nonlinear and nonstationary data. Applying the 3D-EMD method [42] to the fMRI map extracts maps from high to low spatial frequencies. The 3D-EMD method, 1) computes the extrema maps of a volume using a 3D-window (a cube) with the size of 3 voxels that specifies the maximum/minimum values strictly higher/lower compared to their adjacent. 2) The size of the extrema filters is then computed based on the maximum Euclidean distance between the extrema points to be used in making extrema envelopes and their smoothness (Adaptive window size). 3) The smoothed extrema envelopes are then calculated using the 3D-window with the size of the computed extrema filters. 4) The mean envelope is then made by averaging the smoothed extrema envelopes. 5) The mean envelope is subtracted from the original volume to extract the first 3D-IMF or SIMF. 6) Finally, if the subtraction of the residual of the volume and the extracted SIMF contains more than two extrema, the process is repeated to extract the next SIMF; otherwise, there is no more SIMF in the residue, and the 3D-EMD process is completed. For a more detailed description of 3D-method, refer to refs. [38, 42].

The first extracted SIMF that contains high spatial frequencies detects more abrupt changes, peaks, and valleys in the data and, therefore, more spatial details in the data. In contrast, the higher SIMFs that contain lower frequencies show smoother variations. Therefore, high-frequency SIMF, by providing more details of the local spatial changes, helps to find sources of local maximum activation with more chance of their activation being captured by the EEG electrodes. We hypothesize that source-localizing EEG bands, constrained by the EMD-based [40, 41] fMRI’s high-spatial-frequency map, provide more details with higher spatial accuracy about the activity of the sources. It primarily benefits the localization of deep sources, as they must have a relatively high intensity to pass different head layers and get to the electrodes on the head surface [5, 10]. Moreover, in patients with seizures, regions of higher-than-normal blood flow (fMRI intensity maximums) during a seizure may indicate where the seizure occurs [43]. Therefore, using the high SIMFs could assist in specifying a seizure starting point more accurately. Consequently, it improves the clinical interpretation of EEG signals and provides better diagnosis for different brain diseases.

Specifically, we add a weight coefficient computed from the map of the fMRI’s high-frequency SIMFs to the EEG lead-field matrix. Weights with higher values show the locations with a higher probability of being the sources of the recorded EEG signals and vice versa [7]. The proposed method can be used as an accurate neuroimaging technique to unveil the brain sources’ activity with high spatiotemporal resolution. We validate our approach using simulating EEG signals and the corresponding fMRI activation map. We evaluate how different activation intensities and distributions of an fMRI active area affect the source localization results of fMRI- and SIMF-informed EEG inverse modeling. We use measurements such as dipole localization error (DLE), Spatial Dispersion, and the Accuracy to validate the proposed method and compare its results to the fMRI-informed EEG inverse modeling.

## 2. Method

### 2.1 Simulating EEG signals

We simulated EEG signals from two randomly selected sparse connected sources with frequencies of ≈5 Hz and sensor-level SNR of -4 dB for 90 subjects as follows:

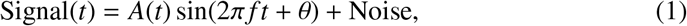

where *A* and *f* are the amplitude and the frequency of the signal, *t* is the time, and θ is the starting phase of the sources’signals. The simulated source signal, is projected to the sensor space using the forward model. 128 scalp electrodes based on the 10 − 20 system and the volume-based BEM model [44] comprising of 13439 vertices (sources) were used to compute the lead field or gain matrix and the forward model. Thus, the sources’ signals are multiplied by the gain matrix to forward project the sources’ signals to the sensors. The noise (usually pink noise, narrow band, or broad band noises) is then randomly added to each of the sensors’ signals to yield the specified SNR (the ratio of signal power to noise power). We simulated sensor noises as the narrow band noises with frequencies in the range of EEG signal’s frequencies.

Signal arises from each source might have a broad range of frequencies. Scalp signals at the EEG sensors are the combination of the captured signals from all the brain sources. Consequently, they contain various frequencies arising from different mechanisms underlying the neural activity [45]. It has been shown that focusing on EEG frequency bands separately and performing source localization for a single EEG band leads to more accurate results than the broadband EEG signal [46]. Accordingly, in this study, we arbitrarily focused on a narrow frequency EEG signal in the frequency range of theta band.

A standard MRI image with size 197 × 233 × 189 was used to make the head model [47–49], and volume-based registration of the EEG sources to be accorded with the fMRI activation map. In this simulation, there was a correlation coefficient of 0.9 and 5 ms delays between the sources’ time courses. We used SIMMEEG [49], a Matlab toolbox, to simulate the EEG signals.

### 2.2 Deriving priors from fMRI

We considered three active areas in the fMRI map. Two corresponded to the two EEG active sources, while the third one had no correspondence in the EEG activation map. For creating the simulated fMRI activation map, fMRI values were drawn randomly from a Gaussian distribution and assigned within a corresponding spatial neighbourhood of each simulated EEG source. We did not designate specific locations for the high-intensity voxels in the active areas; however, we put random values with a higher mean value for closer distances and with a lower mean value for farther distances (Gaussian model). Fig. 1 represents the locations of EEG sources, S1 and S2, which are also the center of the simulated fMRI active regions.

**Figure 1:**
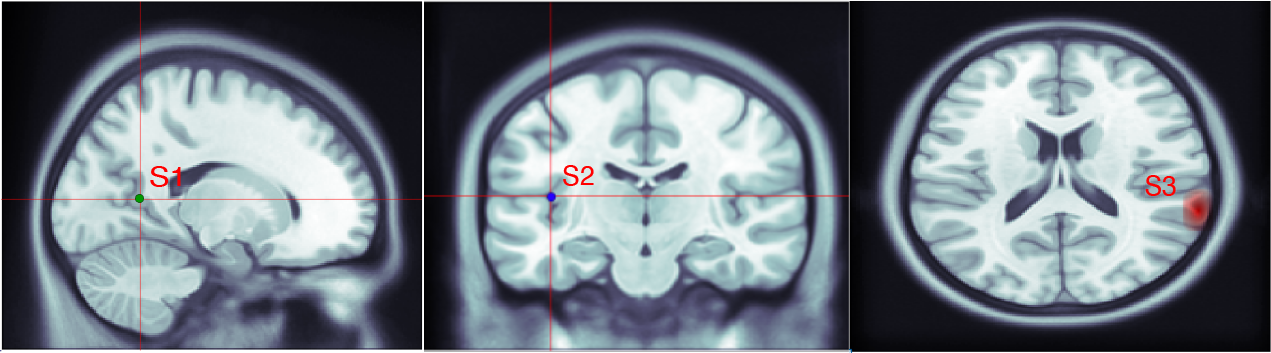
Locations of the simulated sources, S1, S2, and S3, are shown from left to right, respectively. Simulated EEG sources’ subject-based coordinates are S1 = [82, 77, 82] and S2 = [57, 117, 84]. fMRI active regions are simulated in their spatial neighbourhood and comprise deep brain locations in the limbic lobe and posterior cingulate for S1, and sub-lobar, and insula, regions for S2. The fMRI extra source S3 is not related to any EEG source and comprises voxels in the temporal lobe and superior temporal areas.

The third active area in the fMRI activation map was simulated like the other two fMRI active regions, although it does not correspond to neuronal electrical activity and is a representation of other processes in the brain, such as the maintenance of membrane potentials and neurotransmitter release and uptake. This is because there might be places detected as active in the fMRI map but not detectable in EEG recordings. These sources are referred to as “fMRI extra sources” [7, 8, 31]. S3 in Fig. 1 demonstrates the location of the third activated area, the fMRI extra source, in the simulated fMRI map. We change the mean value of the third source’s activation intensity with the same variance as the two other active areas to evaluate the performance of the proposed method for different fMRI activation distributions.

The fMRI-based constraints are added to the EEG inverse problem as a weight coefficient multiplied by the gain matrix. Thus, based on the fMRI activation map, a diagonal location-weighting matrix is computed with a size corresponding to the EEG head model and then is used in the forward model. To compute fMRI’s high spatial-frequency-based weights for EEG inverse modeling, we used the spatial EMD or 3D-EMD method to decompose the fMRI map into its spatial components [42].

Accordingly, the fMRI map is decomposed as follows:

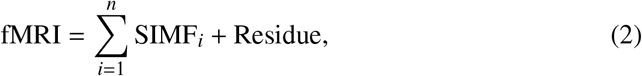

where the first SIMF contains the highest changes of the fMRI activation map while the later SIMFs are smoother maps of spatial changes; the Residue represents the voxels’ intensity trend in the fMRI map.

### 2.2 SIMF-informed EEG Inverse Modeling

Electrical sources *J* of the EEG signals are localized in the brain via inverse modeling. Among different inverse modeling methods, we use the distributed dipole model in this paper. In the distributed dipole model, the magnitude of the sources located on a fixed predefined grid and possibly with fixed orientations are computed using a linear estimation based on Poisson’s equation [17, 29, 50]. The linear model is solved using the L2 or Tikhonov regularization by adding a regularization cost function [9, 13, 15–17, 29, 50]. Thus, sources matrix *J* (the rows are the primary current density in space, and the columns represent time) are computed as follows:

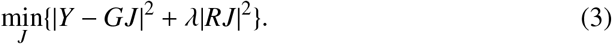

The minimization of the equation (3) yields sources *J* as follows:

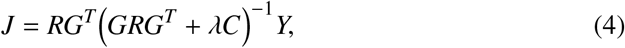

where *G* is the gain matrix of size *N*_e_×3*N*_s_ (*N*_e_ and *N*_s_ are the number of electrodes and sources, respectively). *λ* is the regularization parameter, *C* is the noise covariance matrix, and *Y* is the measured EEG signal at the scalp; its rows and columns represent sensors and time, respectively. The covariance matrix *R* incorporates different constraints on the inverse solution and is based on prior information on the sources activities [9, 29]. In the minimum norm method, *R* is assumed as an identity matrix, *I*, but in the wMNE, it is a diagonal matrix with elements associated with the norm of gain matrix columns, or in LORETA is the spatial Laplacian operator combined with the wMNE gain matrix normalization [9, 21, 26, 29]. For adding the activation information from fMRI to EEG source localization, *R* is considered as a diagonal weighting matrix whose weighting parameters are computed based on the fMRI activation map [6, 9, 13, 15–17, 21, 29, 50].

We used the hig-frequency SIMF for making the diagonal SIMF-induced weighting matrix as the prior variances for EEG sources and to inform the EEG inverse problem. High-frequency SIMF specifies voxels with high intensities relative to their neighbors voxels in the fMRI map, which are more likely to be caused by neuronal electrical activity and firing [32]. We then can apply the intensity threshold to the specified voxels on the fMRI map to exclude voxels of intensity unlikely to be captured by the EEG electrodes and avoid wrong weighting the EEG inverse model. In the simulation, we arbitrarily excluded voxels of intensity less than half of the peak voxel’s intensity. It should be noted that, for real data, the threshold determining enough intensity for a voxel to be recorded on the scalp depends on the location of the activation and its distance to the head surface. Thus, after identifying the locally highly activated voxels using the high-frequency SIMF map, the intensity threshold is applied to the specified voxels on the fMRI map.

The Brainstorm Matlab toolbox [51] was used to solve the EEG inverse problem and localize EEG sources constrained by the fMRI- and high-frequency SIMF-induced weights based on the wMNE method. We used the volume-based BEM model head model, the same volume head model, to avoid errors and mislocalization caused by different head models during inverse and forward modeling. We then registered the EEG head model to the MNI space [52, 53], the same space as the fMRI, to align the fMRI voxel’s coordinates with the head vertices using the nearest neighbor and add the fMRI-based weights to their associated gain matrix’s elements. The same weight was added to the three columns of the gain matrix corresponding to each dipole source.

In informed inverse modeling, large values for the weights derived from the fMRI and its SIMFs activation map indicate more likely active locations. In contrast, small values indicate locations that are less likely to be active in EEG. Furthermore, to avoid spatially biased source localization by using the fMRI map as hard constraints, we put a weight of 0.1 instead of zero as a weight for the gain matrix of the EEG inverse model in not-activated areas in the fMRI map. The off-diagonal elements of *R* were set to zero [8].

Using the Brainstorm, the noise covariance was estimated from the recording data, in which data are baseline corrected by splitting into blocks and subtracting the mean value of each sensor’s data from each block. We examined how different distributions of the sources’ activation affect the source localization result. Fig. 2 presents a schematic of applying the EMD-based-spatiotemporal fMRI constraints to the EEG source imaging.

**Figure 2.**
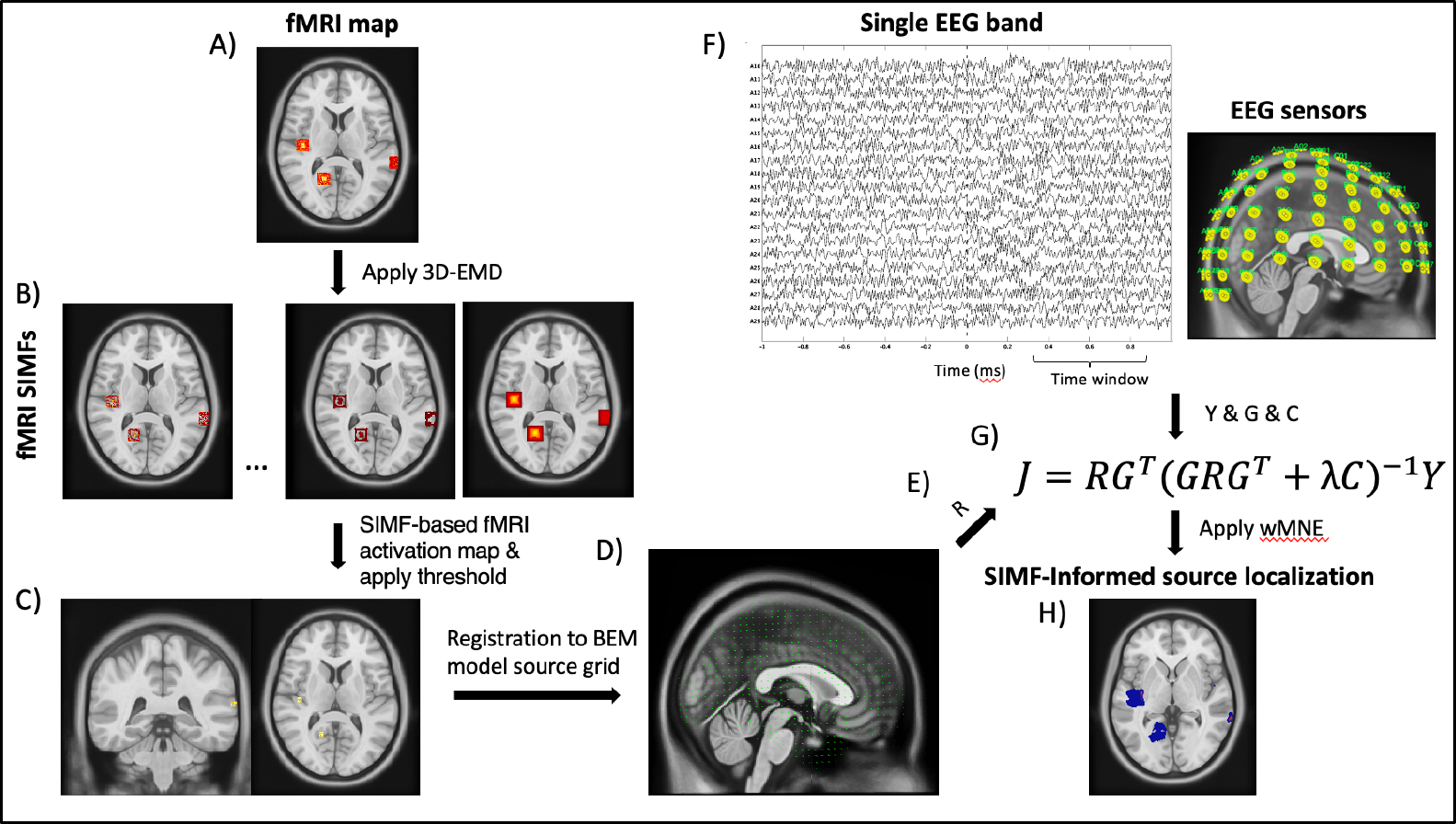
Schematic of SIMF-informed EEG source localization. High-frequency spatial components extracted from the fMRI map and the EEG signals recorded at the same condition are used for finding the activated sources with high spatial and temporal resolution. We start with computing the weighting matrix from the fMRI activation map. A) The 3D-EMD method is applied to the fMRI activation map to decompose it into its B) SIMFs. C) The local highly activated voxels are specified using the high frequency SIMFs and the activation threshold is then applied to the high-frequency-based fMRI activation map. D) The resultant fMRI map is registered to the same head model used for EEG inverse modeling, or vise versa. E) Each diagonal element of the weighting matrix *R* is computed according to its correspondence voxel activity in the fMRI activation map registered to the head model of the EEG inverse modeling. We used a weight of 0.1 for the regions that are not activated based on the fMRI map. F) For a single EEG band (*Y*) recorded at EEG sensors, the gain matrix (*G*) and the covariance matrix (*C*) are computed. G) We solve the fMRI-informed-EEG inverse modeling equation by adding the diagonal weighting matrix *R*, computed from high-frequency SIMFs, as a coefficient for the gain matrix *G*. H) The SIMF-informed EEG source localization result is computed.

To evaluate the proposed method and compare its results with the fMRI-informed EEG source localization, we computed DLE, Spatial Dispersion, and Accuracy parameters [18, 20, 54]. DLE is defined as the Euclidean distance between the true or simulated activated source (*r*_*t*_) and the estimated source (*r*_*e*_) and computed as

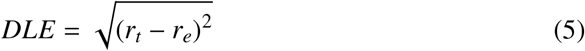

Spatial Dispersion (SD) measures the accuracy of a method in recovering the activated sources within the simulated source space [18, 20, 54]. It is represented by the weighted summation of the distance between estimated and true sources as:

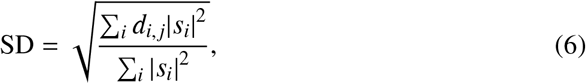

where *s*_*i*_ is the estimated sources activity and *d*_*i, j*_ is a matrix reflecting the DLE value for all sources, the minimum distance between the simulated sources *r*_*t*_ and *i*-th estimated active source [18, 20, 54]. We also computed the accuracy, which evaluates the ability of a method to correctly identify the activated and not activated sources. Accordingly, accuracy is calculated as the number of correct estimations of activation divided by the total number of sources. The Accuracy value is between 0 and 1 [55]. The lower the DlE and SD and the higher the Accuracy, the better the method’s performance in localizing active sources [18, 20, 54, 55].

### 2.4 Real MEG and fMRI data

In this study, task-based MEG data with its corresponding fMRI data from the Human Connectome Project [56] were used. MEG and EEG have similar underlying principles, and an inverse modeling method for either of them applies to both [57]. A brief description of the data is provided below.

#### 2.4.1 MEG motor and working memory tasks data

We used motor and working memory tasks MEG data scanned on a whole head by MAGNES 3600 (4D Neuroimaging, San Diego, CA) [56]. The motor task experimental protocol was adapted from Buckner and colleagues [58, 59]. The Participant was asked to either tap left or right index and thumb fingers or squeeze left or right toe paced by an external visual cue, which was presented in a blocked design. The cue was presented for 3 seconds and the duration of each block of a movement type was 12 seconds. There were also 15-second interleaved resting or fixation blocks per run.

The memory task was designed by Barch and colleagues [60]. During the memory task, participants completed 2-back and 0-back memory tests in two five-minute runs. Participants were presented with four categories of pictures, including places, tools, faces, and body parts as stimuli. Each task block started with a 2.5-second cue specifying the task type. Each run consisted of eight task blocks (10 trials of 2.5 seconds each, for 25 seconds) and four resting fixation blocks. The stimulus at each trial was presented for 2 seconds, followed by an interval time of 0.5 seconds.

Muscle movement signals associated with each scan were measured using the Electromyography electrodes. Electroculography and Electrocardiography electrodes were also used to record heart- and eye movement-related electrophysiological activity. MEG data was preprocessed, including the removal of artefactual-independent components, bad samples, and channels. Data used in this paper were segmented into epochs time-locked to the onset of the Electromyogram signal for motor task data and to the onset of an image that the subject has to match or not with the target image for memory task. The epochs were then categorized into groups of trials representing various events of interest. The average (across trials/runs) event-related fields were then computed at the sensor level.

#### 2.4.2 fMRI motor and working memory tasks data

We used motor and working memory tasks fMRI data of a healthy subject from the Human Connectome Project (HCP) data. Imaging data was recorded by a 3-T Siemens Skyra scanner using a multi-band sequence temporal resolution of 0.72 sec and a 2-mm isotropic spatial resolution. The motor task fMRI data consisted of two 3.5 minutes (284 frames) runs, and the working memory task fMRI data included two 5 minutes (405 frames) runs.

We used fMRI data recorded from the same subject and task [58, 59] as MEG data. fMRI data were minimally preprocessed, e.g., motion correction, distortion correction, and registration to standard space. Further details on fMRI data acquisition, task design, preprocessing and analysing the data can be found in [56, 58–60]. We performed a volume-based GLM analysis for each task to estimate the average effects across runs, using the FSL’s FEAT and the design and setup information of the dataset, e.g., the onset of a stimulus, sequence, and type of each trial.

We applied the proposed method to the fMRI and MEG data associated with the same specific tasks. In this study, we focused on data recorded during the left toe and working memory task, 2-back memory test, to source localize MEG data guided by the fMRI activation map and qualitatively compared the impact of fMRI and its high-frequency spatial component on the accuracy of the source localization.

We source localized the real MEG data like the simulated data using the Brainstorm toolbox and based on the wMNE method. For real MEG data, we used overlapping spheres volume head model and then registered the head model from the subject’s native volume space to the standard MNI space [52, 53] to align the fMRI activation map to the MEG head model’s vertices. We then assigned the fMRI-based weight coefficients (R matrix) for the gain matrix’s elements. We first did not apply the intensity threshold to the high-frequency SIMF-based fMRI activation map to show the effect of SIMF-based weighting compared to the uninformed and fMRI-informed MEG inverse modeling. We then applied a small intensity threshold to the resulting SIMF-based fMRI activation map to avoid the possibility of losing information yet show the effect of the threshold. We removed activations that were lower than 10 percent of the maximum activity for motor task.

## 3. Results

The 3D-EMD method is applied to the fMRI map to decompose it into its SIMFs at each selected time window. Fig. 3, demonstrates the application of the 3D-EMD on the fMRI map used in this study. For illustration purposes, we focused on one of the fMRI active zones to show the result of applying spatial EMD to the data and how the decomposed SIMFs look like. SIMF1 is shown in Fig. 3B and indicates that it can be used to locate the local high-intensity values in the fMRI map and to specify detailed intensity variation in the active area.

**Figure 3.**
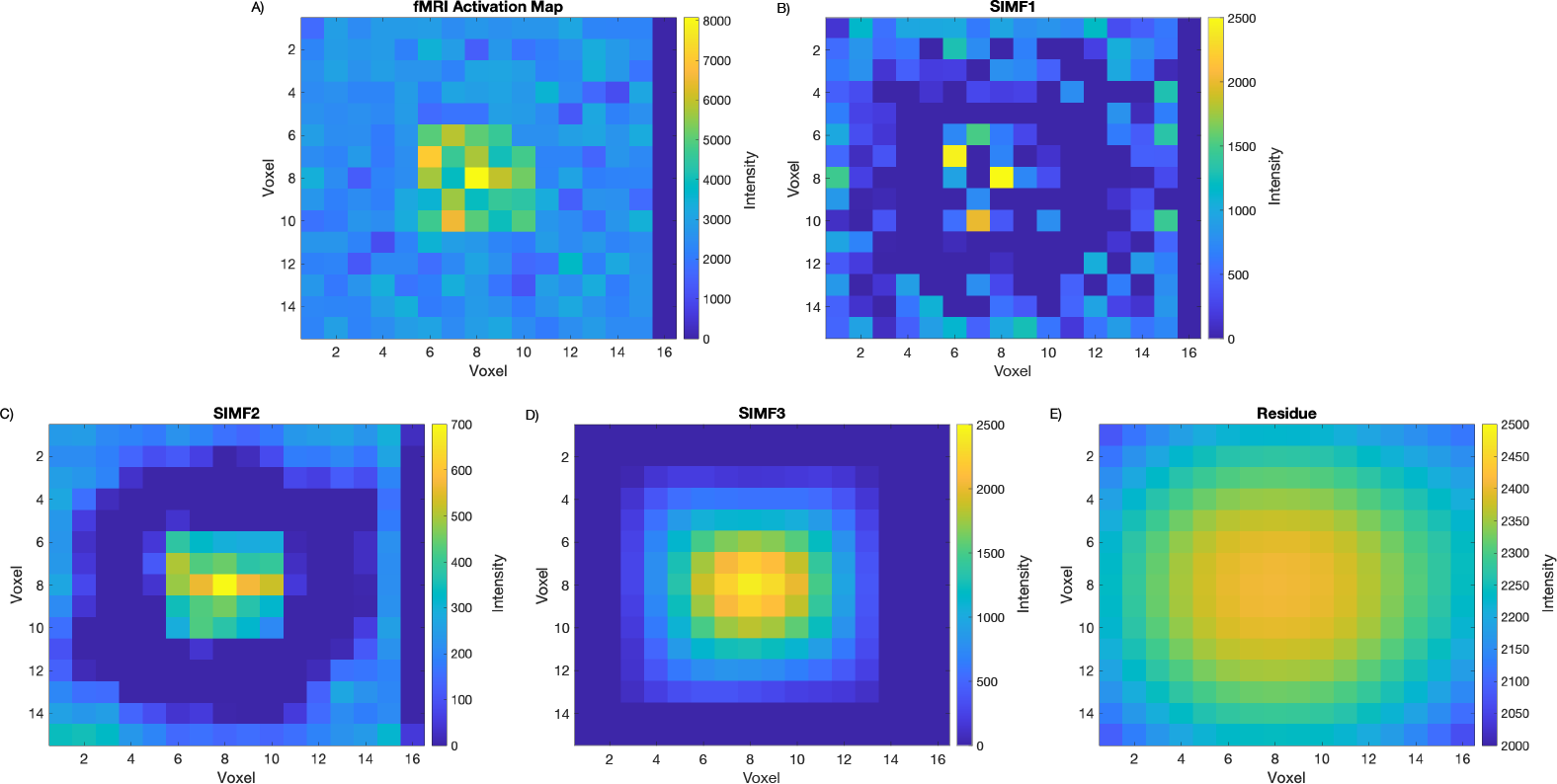
fMRI activation map and its SIMFs around one of the simulated sources. Figures demonstrate A) an fMRI active spot in the brain, B) SIMF1, C) SIMF2, D) SIMF3, and E) Residue, applying the 3D-EMD method. SIMF1 specifies the local maxima of fMRI intensities, and the residue presents the intensity trend of the fMRI activation map’s voxels.

We compare the results of EEG source localization guided by the whole fMRI signal and the high spatial frequency of fMRI for simulated source configurations with different intensity distribution of the third active area. µ is the mean magnitude of the fMRI map at two of the active areas, and the mean magnitude of the third region is changed with respect to them. In this study, we consider different intensity distributions of the fMRI extra source as it is farther from the true EEG sources and variation in the estimated sources in this region makes more distinguishable changes in validation measurement sensitive to the distance between the true and the estimated sources such as the DLE and the Spatial Dispersion compared to the cases where the distribution at the true sources’ area differ.

Table 1 shows the results of the mean DLE over the 90 subjects to compare the fMRI- and SIMF-based EEG source localization for the same source configuration. It indicates that DLE values remain roughly the same with changing the intensity distribution (changing the mean intensity) of the sources for fMRI-informed EEG-source localization. However, for SIMF-informed EEG source localization, the DLE values decrease by decreasing the mean intensity of sources. The lower DLE value represents the better performance of the SIMF-informed EEG source localization in specifying active sources more accurately and with lower localization errors.

**Table 1:**
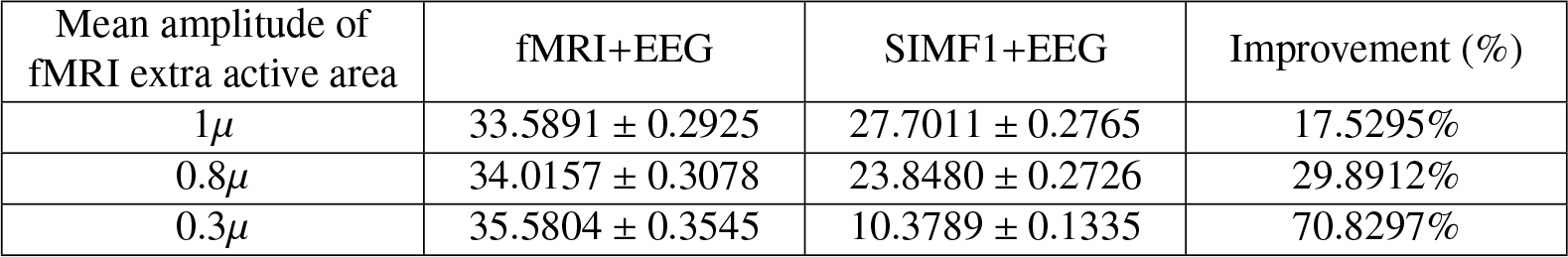
Mean DLE values ± standard deviation for different fMRI extra active area’s amplitude when the whole fMRI signal and high spatial frequencies of fMRI are considered as prior for EEG source localization with p-value < 0.05. Lower DLE means better source localization. The level of the DLE improvement comparing the SIMF-based with the fMRI-based EEG source localization is represented in the third column of the table.

| Mean amplitude of fMRI extra active area | fMRI+EEG | SIMF1+EEG | Improvement (%) |
| --- | --- | --- | --- |
| $1\mu$ | $33.5891 \pm 0.2925$ | $27.7011 \pm 0.2765$ | 17.5295% |
| $0.8\mu$ | $34.0157 \pm 0.3078$ | $23.8480 \pm 0.2726$ | 29.8912% |
| $0.3\mu$ | $35.5804 \pm 0.3545$ | $10.3789 \pm 0.1335$ | 70.8297% |

Table 2 presents the results of Spatial Dispersion for the same comparison as Table 1. For fMRI-informed EEG source localization, the Spatial Dispersion value decreases as the sources’ mean intensity decreases. The lower values of the source intensities in the denominator of the Spatial Dispersion equation make it lower. However, for each source configuration, SIMF-informed EEG source localization shows significantly lower values for the Spatial Dispersion than the fMRI-informed EEG source localization, which represents the better performance of informing EEG source localization by the SIMF rather than the whole fMRI. Table 3 shows the Accuracy results for the same comparisons and source configurations as Table 1 and Table 2. Although substantial improvement is not seen in terms of Accuracy measurements, still, for each source configuration, Accuracy for the SIMF-informed EEG source localization has a higher value than the fMRI-informed EEG source localization.

**Table 2:** Mean spatial Dispersion values ± standard deviation for different fMRI extra active area’s amplitude when the whole fMRI signal and high spatial frequencies of fMRI are considered as prior for EEG source localization with p-value < 0.05. Lower Spatial Dispersion means better source localization. The third column of the table represents the spatial Dispersion improvement percentage, comparing the fMRI and SIMF-based EEG source localization.

| Mean amplitude of fMRI extra active area | fMRI+EEG | SIMF1+EEG | Improvement (%) |
| --- | --- | --- | --- |
| $1\mu$ | $60.1786 \pm 0.6162$ | $58.1358 \pm 0.6354$ | 3.3945% |
| $0.8\mu$ | $61.2021 \pm 0.6160$ | $54.8200 \pm 0.6616$ | 10.4279% |
| $0.3\mu$ | $65.6162 \pm 0.6005$ | $32.9865 \pm 1.1856$ | 49.7281% |

**Table 3:** Mean accuracy values ± standard deviation for different fMRI extra active area’s amplitude when the whole fMRI signal and high spatial frequencies of fMRI are considered as prior for EEG source localization with p-value < 0.05. Accuracy evaluates a method for identifying true activation. A higher value of the Accuracy means better identification of the sources. The third column of the table represents the accuracy improvement percentage, comparing the fMRI and SIMF-based EEG source localization.

| Mean amplitude of fMRI extra active area | fMRI+EEG | SIMF1+EEG | Improvement (%) |
| --- | --- | --- | --- |
| $1\mu$ | $0.8257 \pm 0.0001$ | $0.8459 \pm 0.0011$ | 2.4464% |
| $0.8\mu$ | $0.8273 \pm 0.0010$ | $0.8717 \pm 0.0001$ | 5.3668% |
| $0.3\mu$ | $0.8339 \pm 0.0012$ | $0.8827 \pm 0.0014$ | 5.9409% |

Overall, results of the DLE, Spatial Dispersion, and Accuracy show that informed source localization performs better and is more accurate when the high spatial frequencies of the fMRI activation map are used as prior information instead of the whole fMRI map. By lowering the mean of the fMRI activation intensity but keeping the same variance at the location of the third source with respect to the two other areas, results show lower DLE and Spatial Dispersion and higher Accuracy applying the high-frequency SIMFs.

These results revealed the impact of the distribution of the voxels’ intensities. An active area with high mean intensity has a smoother surface and variation compared to an active area with lower mean intensity and the same variance. Accordingly, for the first case, there might be a case that the whole active area is picked up by the first SIMF. In this case, the SIMF- and fMRI-informed source localization results would be the same. However, the latter case with a lower mean has a relatively sparser and lower number of voxels with high enough SNR, which cause much difference between the activation map derived from high-frequency SIMF and the whole fMRI map. The more locally sparse distribution of high-intensity voxels, the more difference between the SIMF- and fMRI-based EEG source localization results.

Figs. 4 and 5, respectively, show an axial view of the results for fMRI-informed and high-frequency SIMF-informed EEG source localization when the mean intensity of the fMRI extra active area is 0.3 µ. Images from the top left to the bottom right indicate the result of each source localization from 0 to 996.1 ms with 66.5 ms apart. Sources shown in the figures have magnitudes higher than 10 percent of the maximum intensity as a threshold [61].

**Figure 4.**
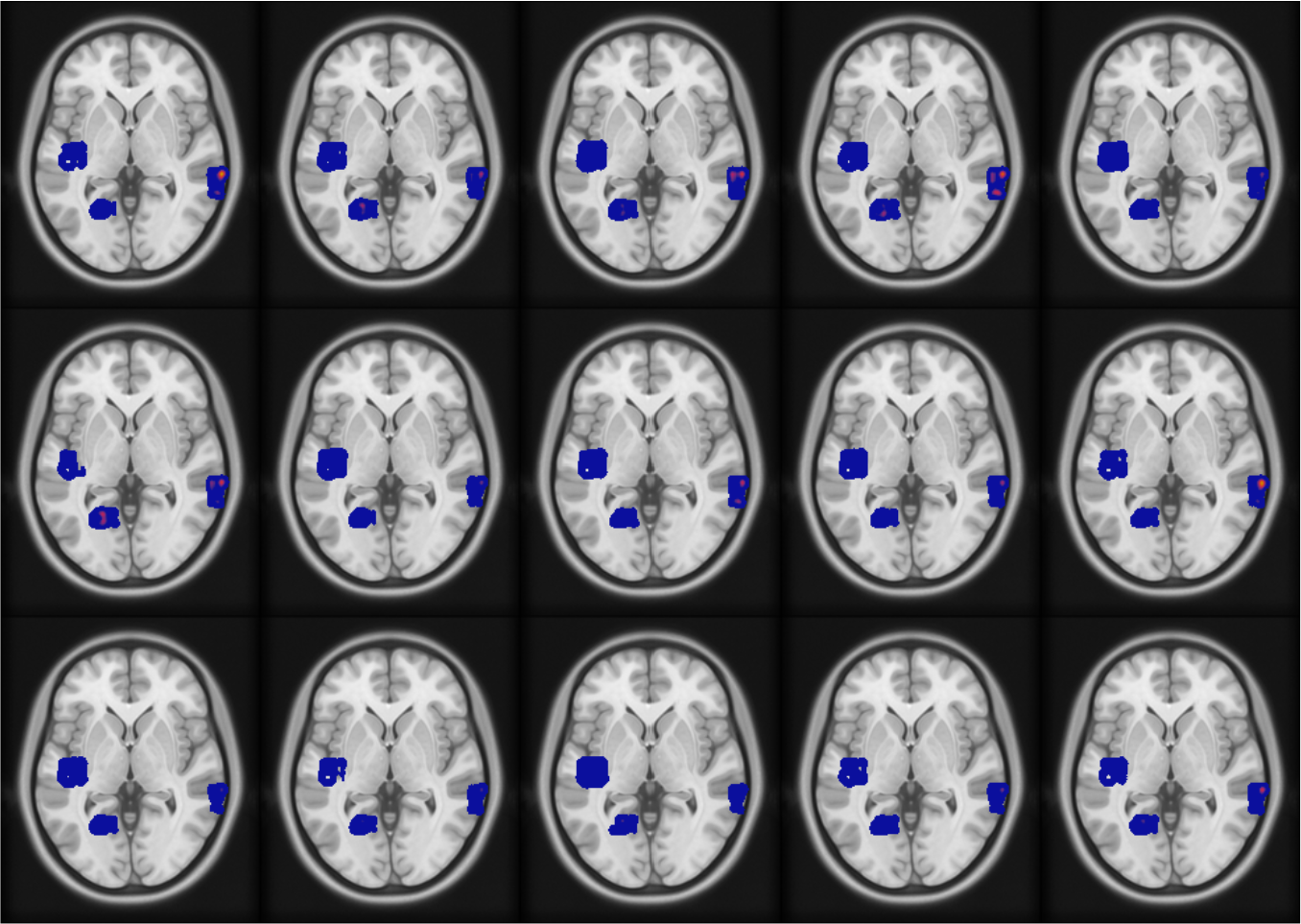
fMRI-informed source localization results when the mean intensity of the fMRI extra active area is 0.3 µ. Axial brain images from top left to bottom right show the sources’ activity from 0 ms to 996.1 ms with 66.5 ms apart.

**Figure 5.**
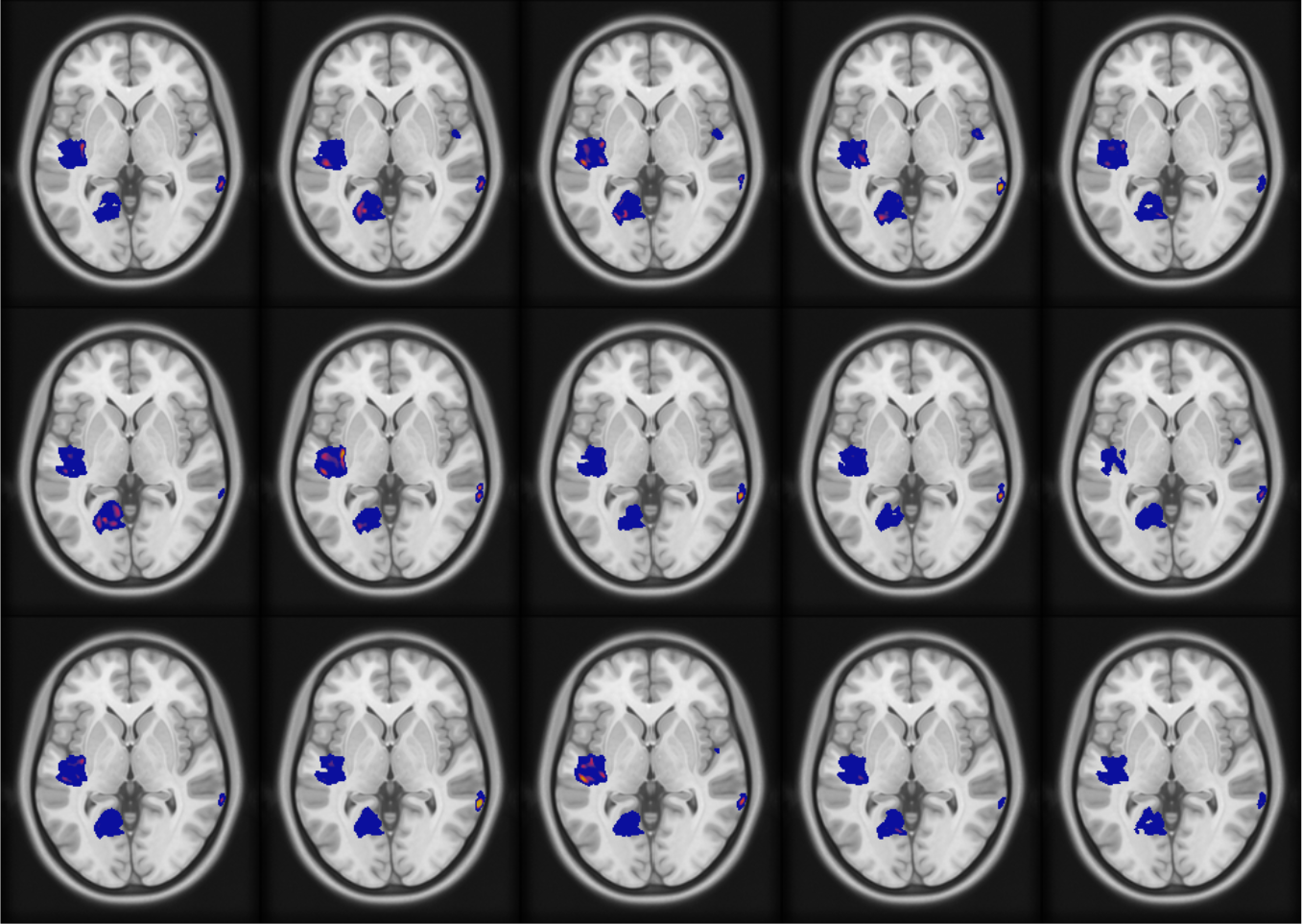
High-SIMF informed source localization results when the mean intensity of the fMRI extra active area is 0.3 µ. Axial brain images from top left to bottom right demonstrate the sources’ activity from 0 ms to 996.1 ms with 66.5 ms apart.

Of note, as the two simulated EEG sources are in deep brain regions, uninformed EEG inverse modeling would show activity in broad brain areas and not in deep confined areas. However, adding fMRI weight to the inverse model helps to find deep activated sources more accurately. Applying SIMF1-induced weights instead of whole fMRI data improves localizing sources’ activity by locating sparser and more accurate local highly active sources with more spatial details and, consequently, higher spatial resolution.

It should be mentioned that we can compute fMRI-based priors for different time window of interest in the EEG data, which makes them time-variant fMRI constraints. It has been shown that EEG inverse modeling informed by static fMRI constraints may cause inaccurate or erroneous results [30].

### 3.1 Experimental result of fMRI informed MEG source localization

We source localized motor task MEG data involving left toe and working memory tasks similar to the simulated data. We anticipated activity in some specific regions in the brain, sensorimotor networks for the motor task (left toe movement) and in prefrontal and parietal brain areas for the working memory task (2-back memory test) according to refs. [56, 58–60, 62].

Figs. 6 and 7 represent the result of MEG data source localization uninformed and informed by their associated fMRI and SIMF data for the left toe and working memory tasks, respectively. Active sources shown in figs. 6 and 7 are of magnitudes higher than 10 percent of the maximum intensity as a threshold [61]. As expected and represented in figs. 6A and 7A, uninformed MEG source localization assigns the activity to wide brain areas and mainly on the cortex surface. Informed source localization, figs. 6B and 7B, specifies active areas more confined and aligned with the designated active areas in [58, 59]. However, SIMF-informed source localization, figs. 6C and 7C, provides more spatially-detailed map of active areas compared to the fMRI-informed source localization. In fig. figs. 6D, we applied a threshold of 0.1 of maximum activation on the high-frequency SIMF-based active areas. We see even a small threshold makes a visible difference as the proposed method by spotting the local maximum activities that are more likely to be associated with the electrical activity, filtering out the BOLD signal extension effects.

**Figure 6.**
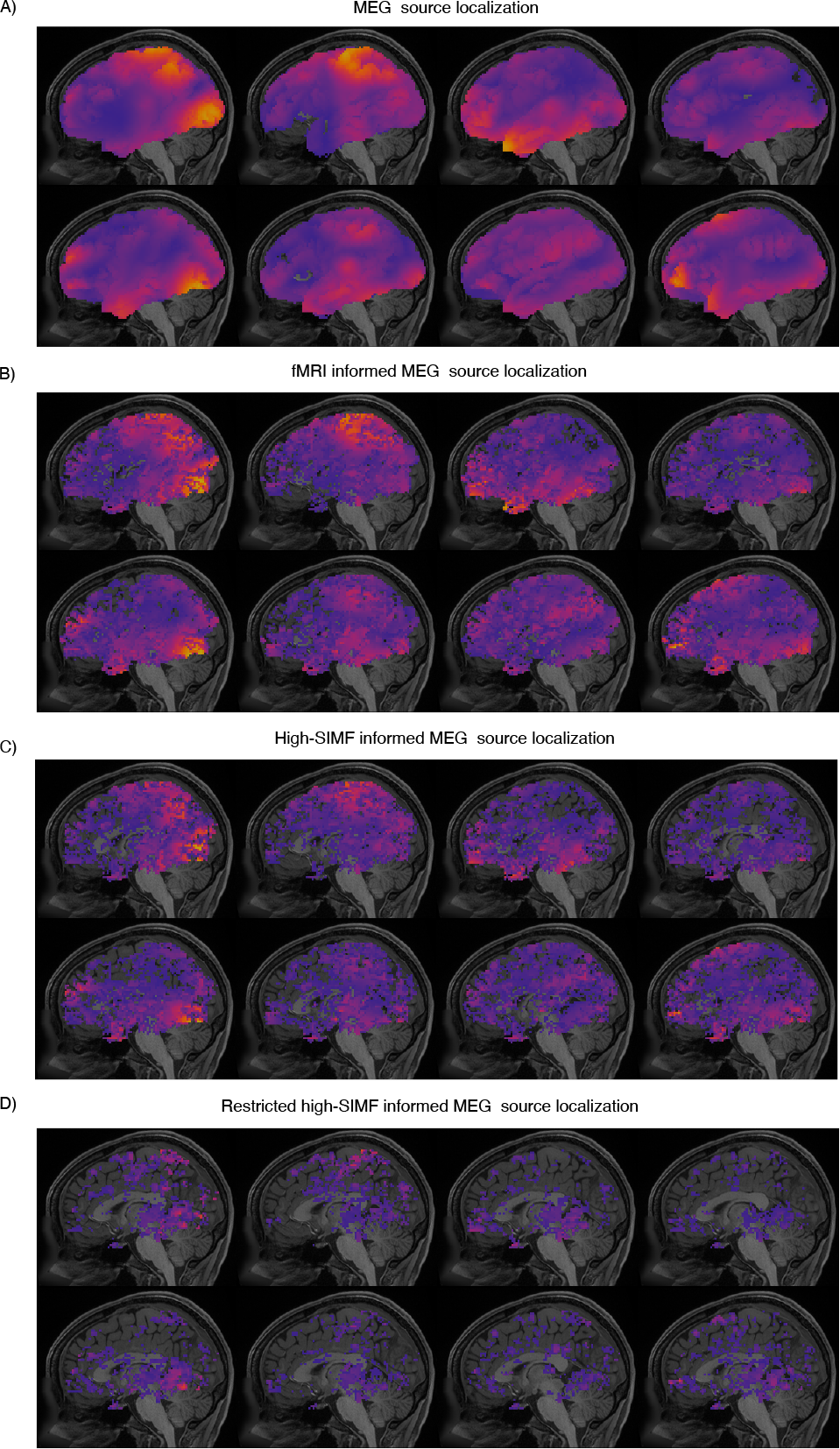
A) Uninformed, B) fMRI- and C)SIMF-informed MEG data source localization for the left toe movements. D) Restricted SIMF-informed MEG source localization, in which voxels with intensities lower than ten percent of the maximum activity on the fMRI map are removed from the weighting matrix added to the MEG inverse modeling. Uninformed source localization results show activation on broad brain areas and more on the cortex surface area. Active areas in fMRI- and SIMF-informed source localization are more spatially specified and in accordance with the expected activity shown in refs. [58, 59]. Active areas localized by the guide of SIMF are more spatially detailed, even without applying any threshold on the fMRI activation map. Figure (D) shows the source localization result when a small threshold (10 percent of the maximum activation intensity) is applied on the SIMF-based activation map and weighting matrix. Brain images from top left to bottom right, for each part of A to D, represent source localization results with ≈125 ms apart.

**Figure 7.**
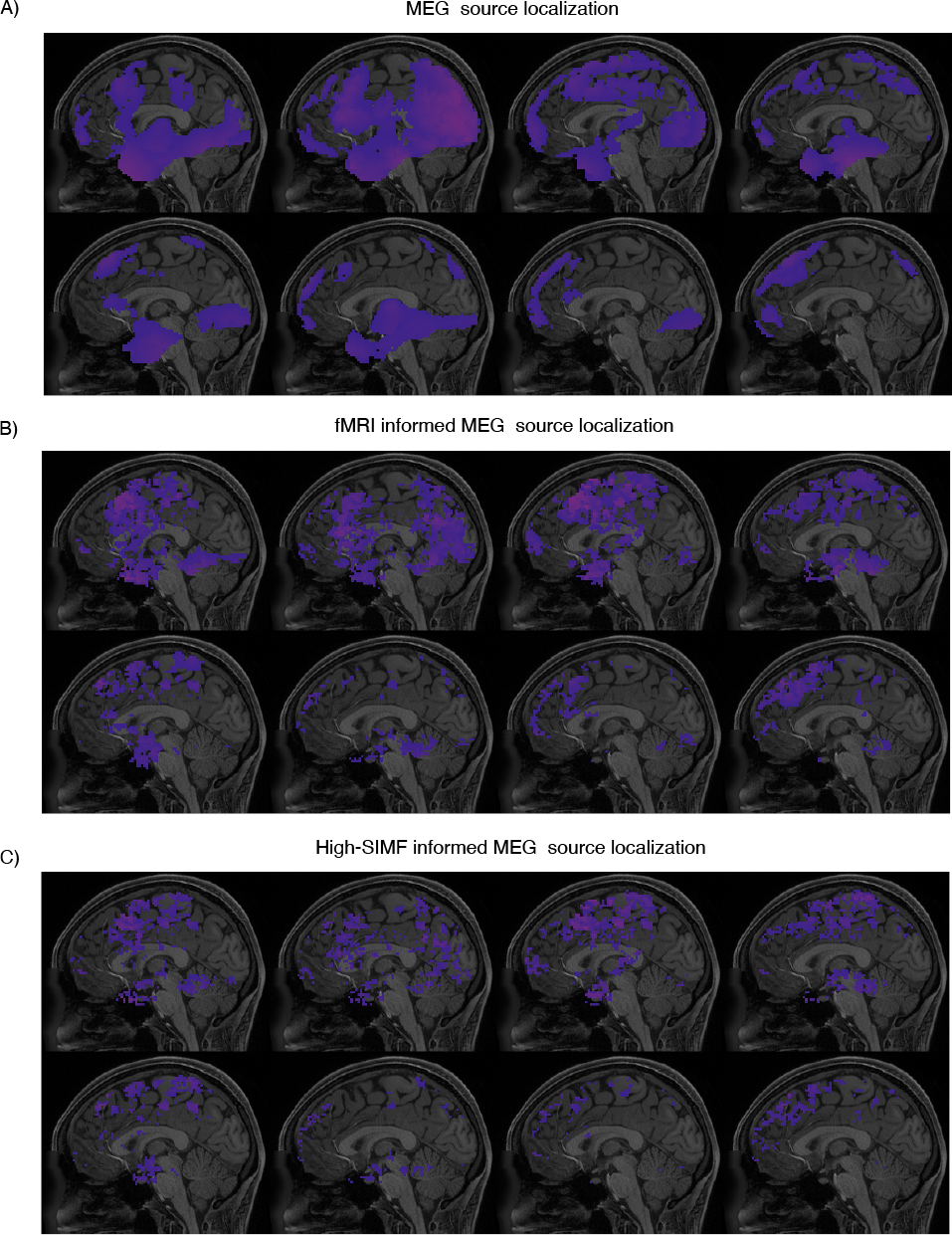
MEG data source localization, A) uninformed, and B, C) informed by their associated fMRI and SIMF data, for the working memory. Informed source localization methods could localize more spatially specified areas and in accordance with the expected activity shown in refs. [58, 59] compare to the uninformed source localization. According to the source localization results shown in figure (C), SIMF-informed MEG source localization leads to more spatially detailed activated areas. The source localization results at each part of A to D, from top left to bottom right, are ≈125 ms apart.

## 4. Discussion

In this paper, we show that using the high spatial components (SIMF) of a fMRI map as a spatial constraint for the EEG source localization improves the accuracy of specifying active sources. This approach is effective for localizing deep brain active sources, which their activities play a pivotal role in central nervous system functioning. Due to the different head-layer conductivities and the distance, electrical potential from deep sources can be captured by EEG electrodes on the scalp if their power is high enough. The larger the depth is, the weaker the electrical signals recorded by the sensors on the scalp. Applying the data-driven 3D-EMD method, fMRI data recorded at the same condition as EEG recording is decomposed into its spatial components (SIMFs). Information from high-frequency SIMF that spot voxels with the local high-activation is added to the EEG inverse model as a weight to localize more confined and precise active sources in all brain areas (deep and surface areas).

Specifically, EEG source localization methods mostly fail to localize deep brain sources [1, 10, 18–20]. They relate the recorded potential to the low-intensity sources on the brain surface instead of high-intensity sources in the deep regions like the insula and thalamus [1, 10, 18–20]. EEG sources are congruent with the hemodynamic changes measured by fMRI. One widely used method to improve this performance impairment of the EEG inverse modeling is using the fMRI map data, which is of high spatial resolution as priors for constraining EEG source space [11, 63, 64]. Thus, several studies have used EEG and fMRI data recorded during the same task or simultaneously in a resting state to yield data with high spatial and temporal resolution [6–9, 29, 30, 39].

However, fMRI-based constraints on EEG source localization may cause spurious results, as all the activities shown on the fMRI map may not captured by EEG sensors. Activities on the fMRI map might not be generated by neuronal firing or not have enough SNR (especially problematic for active deep brain regions) to be recorded by EEG sensors. The first issue might be caused by the spatial extent of the BOLD signals, which appear as voxels with low SNR around highly activated voxels. The second issue is caused by the requirement that neural activity in deep regions, mapped in fMRI, must have an intensity higher than a threshold, depending on the different brain layers’ conductivity, to be recorded by the scalp electrodes.

Subcortical regions are mediators for the brain networks’ communication as they are anatomically connected to an extensive part of cortical regions. Deep regions process primitive functions such as sleep, wakefulness, consciousness, learning, and memory [1–4]. It has been shown in Refs. [2, 65] subcortical structures, such as the amygdala and thalamus, and some parts of the hippocampus, such as the dentate gyrus, have relatively high source density compared to other regions in the neocortex. Thus, a small active volume in these regions produce a high-power signal to be detected by the scalp EEG sensors. fMRI’s high-frequency SIMFs could spot more sparse and confined regions. Consequently, a weighted gain matrix based on the high-frequency SIMFs leads to more precise EEG source localization results and, accordingly, higher resolution compared to when the whole fMRI map is used.

We compared the SIMF-based EEG inverse modeling performance with the fMRI-based EEG inverse modeling. Results demonstrated that using SIMF as a prior constraint makes the EEG inverse modeling more accurate and localizes sources sparser compared to other methods. More accurate source localization means the proposed method provides higher spatial resolution data of neural activity [66].

The results of applying different methods on real MEG and fMRI data for motor and working memory tasks could not be evaluated analytically. However, source localization results applying the SIMF-based method intuitively showed a more localized and spatially detailed map of sources aligned with the expected active areas for the tasks compared to the fMRI-informed and uninformed MEG source localization. Moreover, results showed improvement in source localization even without applying any intensity threshold on the fMRI map and just by removing the effect of the spatial extent of the BOLD signal using the high-frequency SIMFs.

In simulation, we considered the fMRI-extra source in our computations for the EEG inverse modeling. There is also a case where we might have an EEG-extra or fMRI-invisible source. EEG-extra source is either active for a short time or is from a few neurons that cannot make significant and detectable metabolism changes to be appeared in fMRI [67]. It has been shown in Refs. [8, 68] that for including the EEG-extra source, we need to find the proper and high enough value as a coefficient for the fMRI-induced weight matrix that lets the EEG-extra source be visible in inverse modeling. Thus, the whole process will be the same as when there is no EEG-extra source, except that after applying the high-spatial-frequency-induced weight, we find a constant value to be multiplied by the whole lead-field matrix that is large enough to balance the weight coefficient matrix.

It is of note that it has been demonstrated that the fMRI BOLD signal is associated with neuronal synchronization across EEG frequency bands (i.e., neurovascular coupling). The powers of the alpha (8-12 Hz) and beta (12-30 Hz) bands correlate negatively with BOLD signal magnitude, while the gamma band (> 30 Hz) power correlates positively with BOLD signal magnitude [45, 69–71]. Thus, after computing the fMRI activation map, the association between the fMRI BOLD signal with neuronal synchronization across EEG frequency bands (i.e., neurovascular coupling) could be used as an extra filter for specifying fMRI-based weights added to the EEG inverse problem.

It should be mentioned that, in studies that aim for localizing sources of a very specific time point of the EEG data (e.g. peak of ERPs), it is not easy to define a corresponding fMRI activation map due to the fMRI low temporal resolution and the temporal mismatch between the EEG and fMRI.

In summary, EEG inverse modeling methods are integrated with fMRI activation maps to recover deep sources that are problematic and challenging as sources located at the brain surface with low-intensity power are privileged. The proposed approach adds information from the EMD-based fMRI’s high spatial frequency components to more accurately inform the EEG inverse modeling and provide dynamic neural activity with higher spatial resolution. Higher spatial and temporal resolution maps of neural activity improve our brain function knowledge. Precise brain activity localization helps us find more effective treatments for brain diseases such as ADHD and epilepsy by providing substantial information about the source of the epileptogenic process and distinguishing patterns of ADHD [1, 43, 72].

